# Repeated cases of allopatric divergence and secondary contact in cattail (*Typha*) speciation

**DOI:** 10.1101/2024.07.02.601742

**Authors:** Alberto Aleman, Aaron B. A. Shafer, Joanna R. Freeland, Marcel E. Dorken

## Abstract

Understanding how speciation unfolds is a central goal of evolutionary biology. Historically, global climatic fluctuations have triggered allopatric diversification. Using genome-wide data, we reconstructed the phylogenetic relationships and demographic histories of five *Typha* species, a plant genus foundational to freshwater ecosystems, with widespread, partially sympatric ranges and at least one widespread, regionally invasive hybrid zone (*Typha* × *glauca*). Molecular clock and demographic analyses indicate that the species in this study diverged in the absence of gene flow, during periods roughly contemporaneous with population bottlenecks and expansions in *Typha*—likely triggered by geoclimatic events—and that this divergence was followed by secondary contact in more recent times. Genomic scans showed no evidence of selection with gene flow driving species differentiation, suggesting a minor role for sympatric ecological speciation underlying divergence. These observations are consistent with expectations for drift-driven allopatric speciation. Allopatric speciation in *T. latifoli*a and *T. angustifolia*, their low genetic differentiation, and the scarcity of ecological divergence in building their reproductive isolation could explain these species’ ability to hybridise.

## Introduction

Speciation can follow two non-mutually exclusive pathways: geographic isolation (“allopatric speciation”) and natural selection under gene flow (“ecological speciation”) (Edwards *et al*., 2020). In allopatry, lineages diverge without gene flow, leading to the independent accumulation of genetic differences—most of which arise through drift—over time (Sobel, 2016). Under gene flow, genetic differences accumulate as divergent selection acts on adaptive traits and the loci underlying them, first restricting gene flow at adaptive loci (Tigano and Friesen, 2016; Elmer, 2019; Stankowski and Ravinet, 2021) and later inhibiting reproduction between lineages (Nosil, 2012). Understanding how both speciation pathways unfold can help us reconstruct species’ evolutionary histories, a central goal of evolutionary biology (Avise, 2000).

Global climatic fluctuations have repeatedly shaped species’ distributions and abundances, creating opportunities for allopatric speciation (Kumar and Kumar, 2018). Major geoclimatic transformations have promoted several drift-driven speciation events (Andersson, 2009; Chacón *et al*., 2019). For instance, aridification during the Mid-Miocene and early Pliocene caused habitat fragmentation, triggering diversification in numerous species (e.g., Shi *et al*., 2013; Manish and Pandit, 2018; Kergoat *et al*., 2018; Ansari *et al*., 2019; Aduse-Poku *et al*., 2021; Dagallier *et al*., 2024; Hühn *et al*., 2024; Adams *et al*., 2025; Yan *et al*., 2025). Such drift-driven speciation events, driven by strong reductions in effective population sizes, can often be identified by linking lineages’ historical bottlenecks to their divergence times (Bock *et al*., 2023).

Cattails (*Typha*) are a widely distributed genus of rhizomatous, perennial, monoecious, self-compatible, wind-pollinated, monocotyledonous flowering plants crucial to wetlands (Grace and Harrison, 1986): they play vital roles in nutrient cycling, preventing erosion, maintaining stable water levels, and providing food and shelter for vertebrates and invertebrates (reviewed in Bansal et al., 2019). The three most widespread *Typha* spp. (*T. angustifolia*, *T. domingensis*, and *T. latifolia*) have extensive areas of sympatry and, in at least some regions, can hybridise with one another (Smith, 1967). Notably, *T. angustifolia* and *T. latifolia* hybridise to form *T.* × *glauca,* which in North America occupies a widespread and expanding hybrid zone (Joyee *et al*., 2024); this hybrid is often considered invasive because it forms dense stands, alters habitats, and outcompetes and displaces native taxa (reviewed in Bansal *et al*., 2019; Freeland and Dorken, 2026).

Historical distributions of cattail species are unknown; however, since hybridisation often results from allopatric divergence followed by secondary contact (Sobel, 2016; e.g., Maguilla *et al*., 2017; Schield *et al*., 2019; Arteaga *et al*., 2020; Kessler *et al*., 2023), we hypothesise that *T. angustifolia* and *T. latifolia* speciated in allopatry, i.e., that they diverged in geographic isolation (under no divergent selection with gene flow), thereby without evolving “barrier” loci (Elmer, 2019; Stankowski and Ravinet, 2021)—before expanding into widespread areas of sympatry, where they can hybridise. To test the hypothesis of allopatric speciation, we reconstructed the phylogenetic relationships and demographic history of *T. angustifolia* and *T. latifolia*, and their sister species, *T. domingensis*, *T. laxmannii*, and *T. shuttleworthii* (Zhou *et al*., 2018), using genomic data from 207 plants sampled across multiple continents (Figure 1).

**Figure 1.**
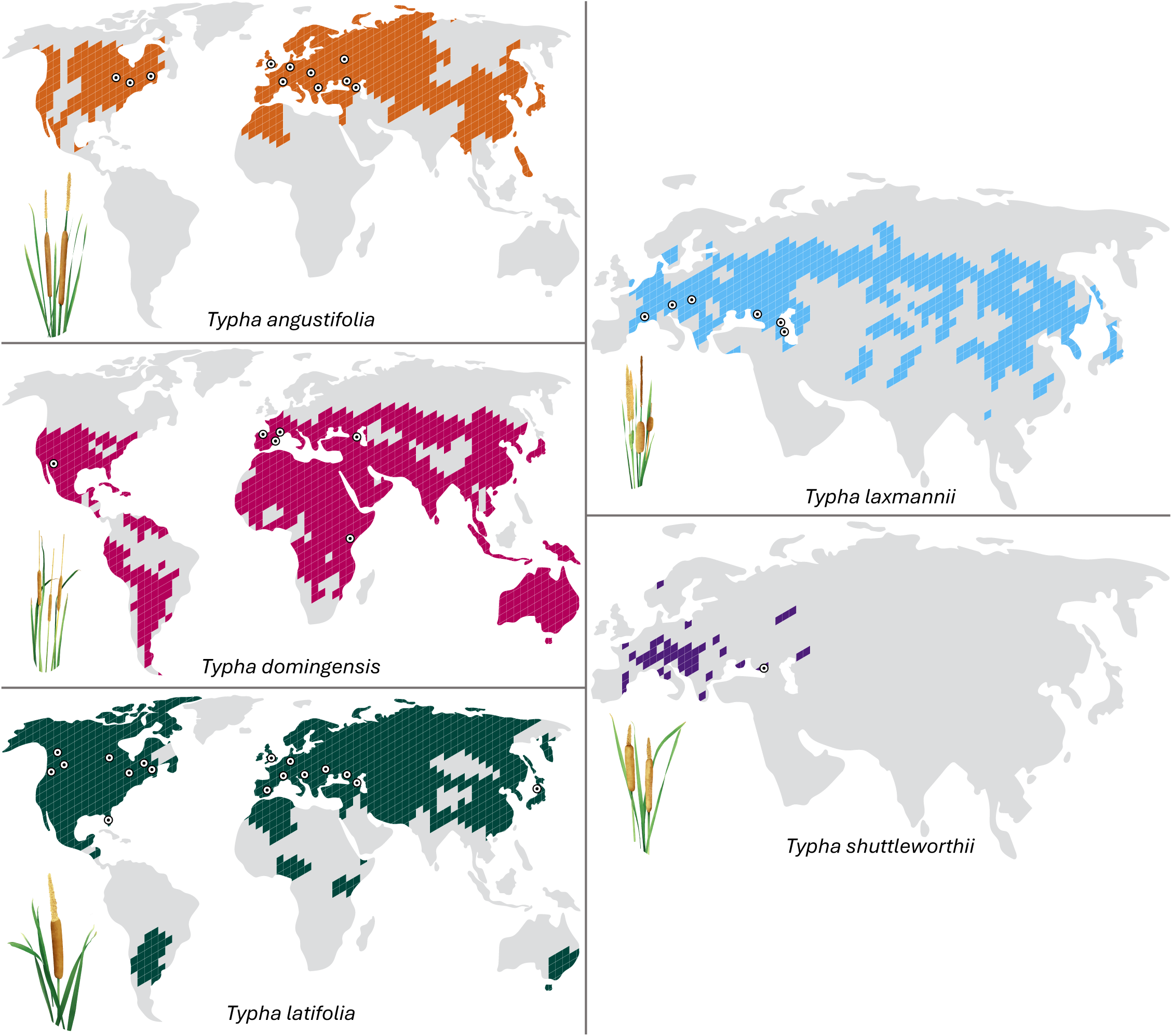
Species distributions (coloured areas) and sample sites in this study (circles) for *Typha angustifolia*, *T. domingensis*, *T. latifolia*, *T. laxmannii*, and *T. shuttleworthii*. Distributions from Ciotir & Freeland (2016) and GBIF. Distributions are based on maps of geopolitical boundaries in areas where species have been recorded and do not necessarily reflect geographic coverage. Sample sizes are not shown graphically.

## Materials and Methods

### Sampling, DNA extraction, and sequencing

Samples were obtained from Aleman *et al*. (2024) and supplemented with additional collections to enhance taxonomic diversity and broaden our sampling range (Figure 1; Supplementary Table S1). Plants were identified using morphological characteristics (Grace & Harrison, 1986; Smith, 1967) and/or by genotyping 3 to 4 microsatellite loci with species-specific alleles (Bhargav et al., 2022; Ciotir et al., 2013, 2017; Ciotir & Freeland, 2016; Pieper et al., 2020; Tisshaw et al., 2020). DNA was extracted as per Pieper *et al*. (2017, 2020) and converted into Nextera XT libraries, following Aleman *et al*. (2024). Paired-end sequencing was conducted on a Novaseq 6000 (126 bp) and a Miseq (151 bp) at The Centre for Applied Genomics (Toronto, Ontario, Canada) for 64 *T. angustifolia*, 25 *T. domingensis*, 104 *T. latifolia*, 11 *T. laxmannii*, and 3 *T. shuttleworthii*.

### Raw data processing

The quality of demultiplexed raw sequences was assessed with FastQC 0.11.9 (Andrews, 2017) and MultiQC 1.14 (Ewels *et al*., 2016). Read pairing and adapter trimming were performed with Trimmomatic 0.39 (Bolger *et al*., 2014), and reads shorter than 100 bp were removed. Cleaned reads were mapped to the *T. latifolia* nuclear (285.11□Mb, GenBank accession JAIOKV000000000.2) and chloroplast (161.57 kb, GenBank accession NC_013823) genomes with BWA 0.7.17 (Li & Durbin, 2009). Mapping statistics were evaluated with SAMtools 1.15.1 (Li *et al*., 2009).

### Genotyping

Genotyping was performed using ANGSD 0.93 (Korneliussen et al., 2014). SNPs were called across all samples (SNP data, henceforth), establishing mapping- and base-quality thresholds of 20 and a minimum p-value of 1e−6. Additionally, chloroplast sequences were called for each sample using ANGSD, with reads mapped to the chloroplast reference and mapping- and base-quality thresholds of 20. Loci with more than 20% missing data and those mapped to the plastome were removed from the SNP data with VCFtools 0.1.16 (Danecek et al., 2011). The absence of clones (multiple ramets from the same genet) was verified by calculating kinship coefficients using Plink 2.0 (Chang et al., 2015) and the SNP data.

### Genetic relationships

Two approaches were used to assess the most likely number of nuclear clusters across all samples and each sample’s membership in these clusters, using SNP data. First, an Admixture 1.3.0 plot (Alexander and Lange, 2011) was run with *K* ranging from 1 to 10, and the optimal number of clusters (*K*) was selected based on the lowest cross-validation error. Because Admixture tends to fit smaller groups as mixtures of multiple groups rather than as distinct clusters (Lawson *et al*., 2018), *T. shuttleworthii* was excluded due to its small sample size (*n* = 3). Then, a principal component analysis (PCA) of all 207 samples was performed using Plink 1.9 (Purcell *et al*., 2007). Complementarily, a phylogenetic tree of all 207 samples was reconstructed from chloroplast sequences using PhyML 3.0 (Guindon *et al*., 2010); model selection was based on the Akaike information criterion (AIC), estimated by SMS (Lefort *et al*., 2017), and PhyML was run under GTR+R with automatic bootstrapping.

### Divergence times

Divergence times among species were reconstructed using cpDNA and a molecular clock in BEAST 1.10.4 (Drummond and Bouckaert, 2015). We selected the chloroplast sequence with the highest data completeness for each species in our dataset, as well as five Bromeliaceae, three *Sparganium*, and two to three additional *Typha* (depending on classification; Supplementary Table S2). Since we observed two lineages in *T. latifolia* (named “Western” and “Eastern”; see *Results*), we included one sequence from each lineage. All plastomes were aligned with MAFFT 7 (Katoh *et al*., 2019). Based on the results of jmodeltest 2.1.10 (Darriba *et al*., 2012; Edler *et al*., 2021), a GTR+G+I model was used. Following Zhou *et al*. (2018), a log-normal prior distribution on the divergence between *Typha* and *Sparganium* was set to 70 Ma (offset = 70, mean = 1.5, SD = 0.5), together with normal distributions at the crown of Bromeliaceae (mean = 19.1, SD = 1) and the crown of the three (mean = 100, SD = 1.7). We ran two independent chains of 110,000,000 steps, sampling every 1,000,000 steps. Runs were joined using LogCombiner, with a 10,000,000-step burn-in per input, and a maximum-credibility tree was produced using TreeAnnotator.

### Demographic histories

Changes in effective population sizes (N_e_) over time for each species were reconstructed using Stairway Plot 2.1.2 (Liu and Fu, 2020). We then modelled the demographic histories of the species in this study using δaδi (Gutenkunst *et al*., 2010) and dadi_pipeline 3.1.7 (Portik *et al*., 2017). Based on their phylogenetic relationships, we reconstructed the demographic histories of *T. angustifolia* relative to *T. domingensis*, and of *T. shuttleworthii* relative to *T. latifolia* (Western and Eastern, since we observed two lineages in this species; see *Results*). We included the models by Portik *et al*. (2017), Barratt *et al*. (2018), and Charles *et al*. (2018) (Supplementary Figures S3–S4), and selected the best models based on the AIC. Folded spectra were generated from the SNP data using easysfs 0.0.1 (Gutenkunst *et al*., 2009), with a 2-year generation time and a µu of 7×10^-9^ mutations per site per generation (Weng *et al*., 2019) in both pipelines.

### Role of selection in species divergence

We examined genetic differentiation and diversity (F_ST_, d_XY_ and π) within and between species following Kessler *et al*. (2023) and Shang *et al*. (2023). Because these metrics require both SNPs and invariant sites, poly- and monomorphic loci were called for all samples using ANGSD, with minimum mapping and base-quality thresholds of 20 and no SNP p-value cutoff. Data mapped to the plastome were removed from this dataset.

The script popgenWindows.py (Martin, 2023) was used to estimate F_ST_ (Weir and Cockerham, 1984), d_XY_ (Nei and Miller, 1990), and π (Nei and Li, 1979) in 5- and 10-kb windows for all species pairs and between the two *T. latifolia* lineages. To ensure data reliability, windows with coverage below 50% were discarded in each comparison. The means for each statistic and the net divergence between taxa (d_a_; Nei and Li, 1979) were calculated in R 4.5.2 (R Core Team, 2022). Outlier windows were then identified and classified according to the models of ecological divergence proposed by Han *et al*. (2017) and Irwin *et al*. (2018): (i) divergent selection under gene flow (high F_ST_ and d_XY_, low π); (ii) divergent selection without gene flow (high F_ST_, intermediate d_XY_, and low π); (iii) background selection (high F_ST_, low d_XY_ and π); and (iv) balancing selection (low F_ST_, high d_XY_ and π). Thresholds were set at the upper or lower 5% for high or low criteria, and at 45–55% for intermediate criteria.

## Results

### Genetic relationships

We assembled ∼20.96M nuclear SNPs across 207 samples and retained 77,207, yielding a mean missingness of 17.1% and a mean depth of 4.82× per sample (Supplementary Table S1). Cross-validation of Admixture, run without *T. shuttleworthii*, indicated *K* = 6 as the most likely number of clusters. Species were discernible at *K* = 5, and *T. latifolia* was assigned to two distinct clusters, broadly identified as “Western” (North America, Western and Central Europe) and “Eastern” (Eastern Europe and Iturup Island, in the Russian Kuril chain). At *K* = 6, a shared component was observed in most samples; since the groupings remained the same as at *K* = 5, we considered *K* = 5 valid (Figure 2; Supplementary Figure S5). The PCA differentiated all species, indicating overlap on PC1 among *T. angustifolia*, *T. domingensis*, and *T. laxmannii*, and on PC2 between *T. domingensis* and *T. laxmannii*, and between *T. latifolia* and *T. shuttleworthii*; however, PC3 differentiated *T. domingensis* from *T. laxmannii*, explaining variance comparable to that of PC2 (15.6% and 17%, respectively) (Figure 2). The two *T. latifolia* lineages were not differentiated by the PCA. The chloroplast phylogeny identified six clades, distinguishing all five species and splitting *T. latifolia* into two clades; consistent with the nuclear genetic structure, these two clades corresponded to the Western and Eastern lineages identified by Admixture (Figure 2). Notably, six nuclear Western samples—two from North America and four from Western and Central Europe—carried the Eastern plastome; no other cytonuclear discordances were observed.

**Figure 2.**
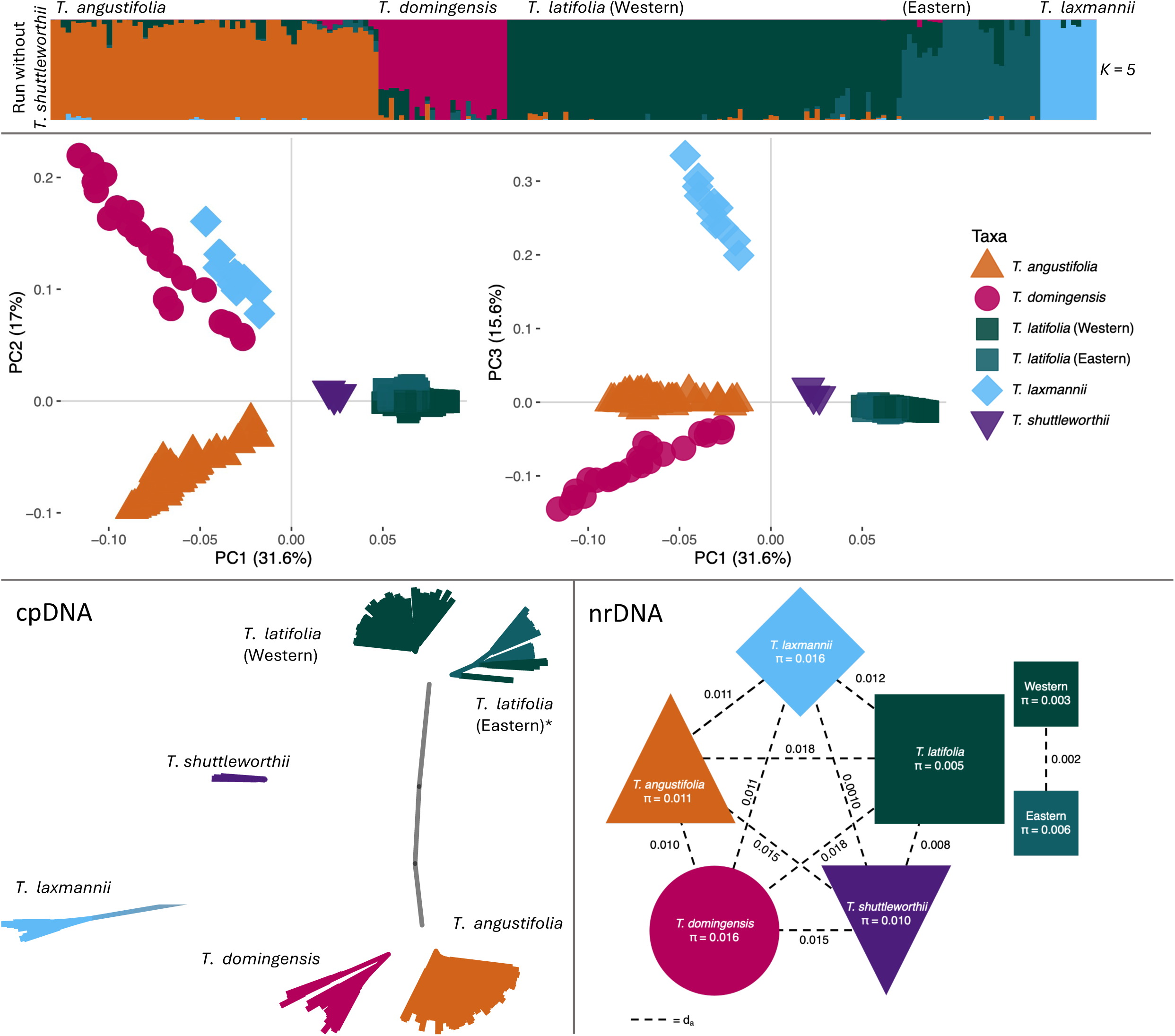
Genetic relationships and species differentiation among five *Typha* species. Top: Admixture plot for *K* = 5; vertical bars represent individuals, and genetic components are shown in different colours. Middle: Principal component analyses. Points represent individuals; colours and shapes denote different taxa. The Admixture plot and PCAs were based on 77,207 nuclear SNPs. Bottom left: Unrooted chloroplast phylogeny (from 161,572 bp); branches represent individuals, and colours indicate species, as labelled. *Six nuclear Western samples carried the Eastern plastome. Bottom right: Nucleotide diversity (π, inside shapes) and net divergence (d_a_, dashed lines) among taxa in this study, from nuclear genomic scans of ∼136.11 Mb. Note: All samples from the two *T. latifolia* lineages were pooled to obtain π and d_a_ for the species.

### Divergence times

The 17 aligned chloroplast sequences used for phylogenetic reconstruction and the molecular clock were 171,915 bp in length. BEAST estimated that the most recent common ancestor (MRCA) of *T. angustifolia* and *T. domingensis* occurred ∼9.9 Ma (95% HPD: 3.5–38.0 Ma), the MRCA of *T. latifolia* and *T. shuttleworthii* occurred ∼10.8 Ma (2.5–33.7 Ma), and the MRCA of the Western and Eastern lineages of *T. latifolia* occurred ∼3.4 Ma (1.5–16.1 Ma). The focal species in this study, including *T. laxmannii*, shared their MRCA ∼41.9 Ma (7.0–61.5 Ma) (Figure 3; Supplementary Figure S6).

**Figure 3.**
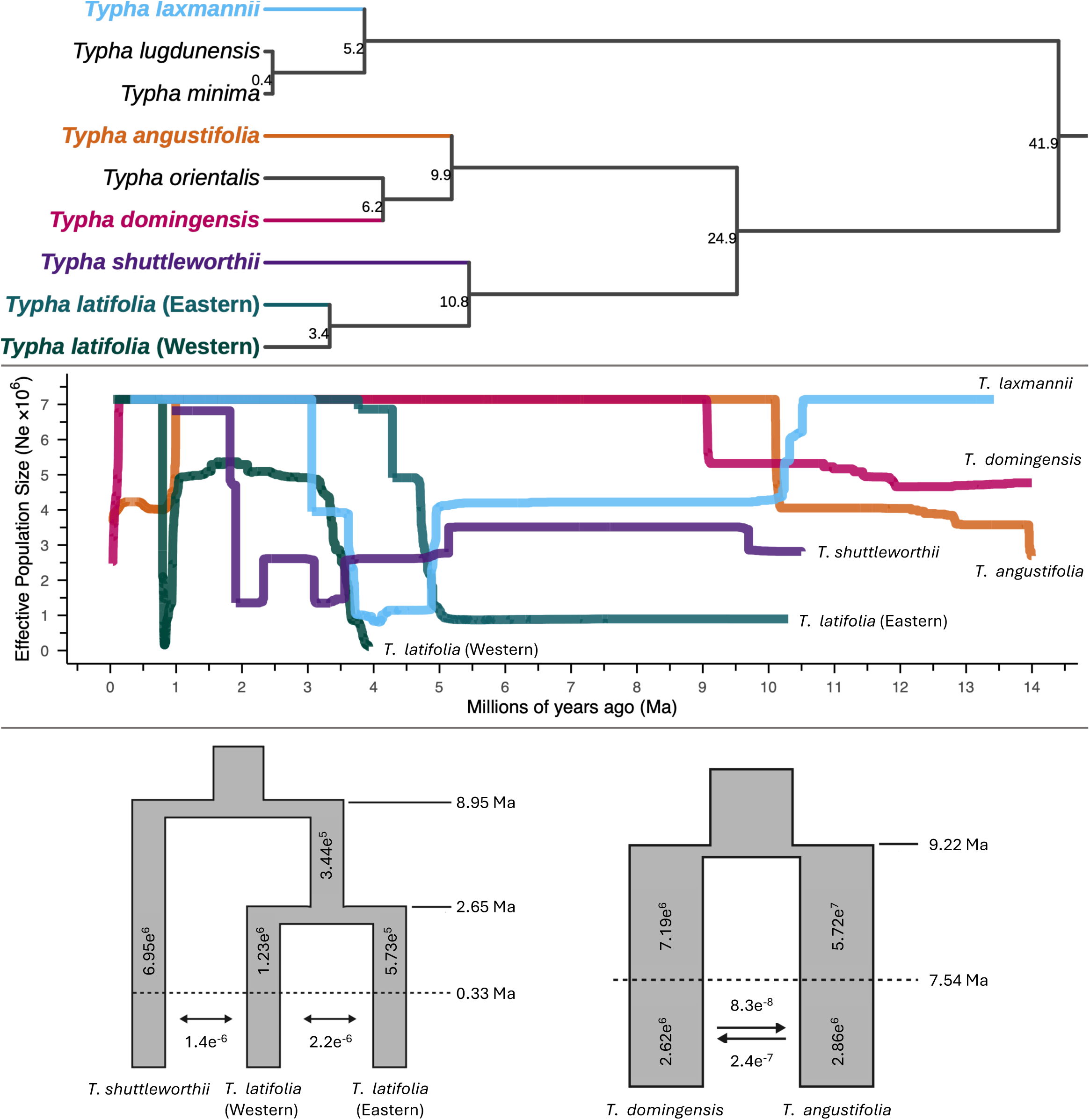
Speciation of *Typha* in this study. Top: Dated phylogeny of eight *Typha* species, inferred from chloroplast sequences (171,915 bp) using BEAST. Note: There is debate over whether *T. lugdunensis* and *T. minima* are distinct species (see GBIF). Middle: Historical N_e_ for five *Typha* spp., reconstructed with Stairway Plot. Bottom: Demographic histories of *T. angustifolia* relative to *T. domingensis*, and of *T. shuttleworthii* relative to *T. latifolia* (Western and Eastern), reconstructed with ∂a∂i. Arrows indicate the number of migrants per generation; vertical numbers indicate N_e_.

### Demographic histories

The reconstruction of N_e_ over time inferred historical bottlenecks and expansions in the lineages leading to the focal species in this study: (i) between ∼10.5 and 9 Ma, the lineage leading to *T. laxmannii* experienced a bottleneck, followed by expansions in the lineages leading to *T. angustifolia*, *T. shuttleworthii*, and *T. domingensis*; and (ii) between ∼5 and 3.5 Ma, the lineages leading to *T. laxmannii* and *T. shuttleworthii* experienced bottlenecks, followed by expansions in the lineages leading to *T. latifolia* (Western and Eastern) (Figure 3).

Modelling of species’ demographic histories suggested that (i) *T. angustifolia* and *T. domingensis* diverged in isolation ∼9.22 Ma and experienced secondary contact ∼7.54 Ma; *T. shuttleworthii* and *T. latifolia* diverged in isolation ∼8.95 Ma, and Western and Eastern *T. latifolia* diverged in isolation ∼2.66 Ma, followed by secondary contact between *T. latifolia* lineages and between *T. shuttleworthii* and Western *T. latifolia* ∼0.33 Ma (Figure 3; Supplementary Table S7–S8). It should be noted that AIC indicated that the optimal model for the demographic history of *T. angustifolia* and *T. domingensis* had no changes in N_e_ over time; given that N_e_ fluctuations are more realistic than a constant size (see *Discussion*), we obtained estimates of population sizes, divergence and secondary contact times, and migration rates from a model that allowed for changes in N_e_.

### Role of selection in species divergence

Based on 47.74% of the genome (136.11 of 285.11 Mb), the net divergence between species was lowest between *T. latifolia* and *T. shuttleworthii* (d_a_ = 0.008) and highest between *T. latifolia* and both *T. angustifolia* and *T. domingensis* (d_a_ = 0.018) (Figure 2). Divergence between Western and Eastern *T. latifolia* (d_a_ = 0.002) was lower than that between species (mean d_a_ = 0.012). Diversity was lowest in *T. latifolia* (π = 0.005) and highest in both *T. domingensis* and *T. laxmannii* (π = 0.016). The genomic landscapes across species pairs showed similar patterns of ecological divergence and revealed an absence of divergent selection under gene flow during species differentiation. Most outlier windows (with exceptionally high or low F_ST_, each 10 Kb) could not be associated with any of the selection types tested (on average, 488 out of 680). Among the outlier windows that could be explained by a model of ecological divergence, balancing selection was the most frequently detected, followed by background selection and divergent selection without gene flow (Table 1). The same patterns were observed with a window size of 5 Kb (Supplementary Table S9).

**Table 1.**
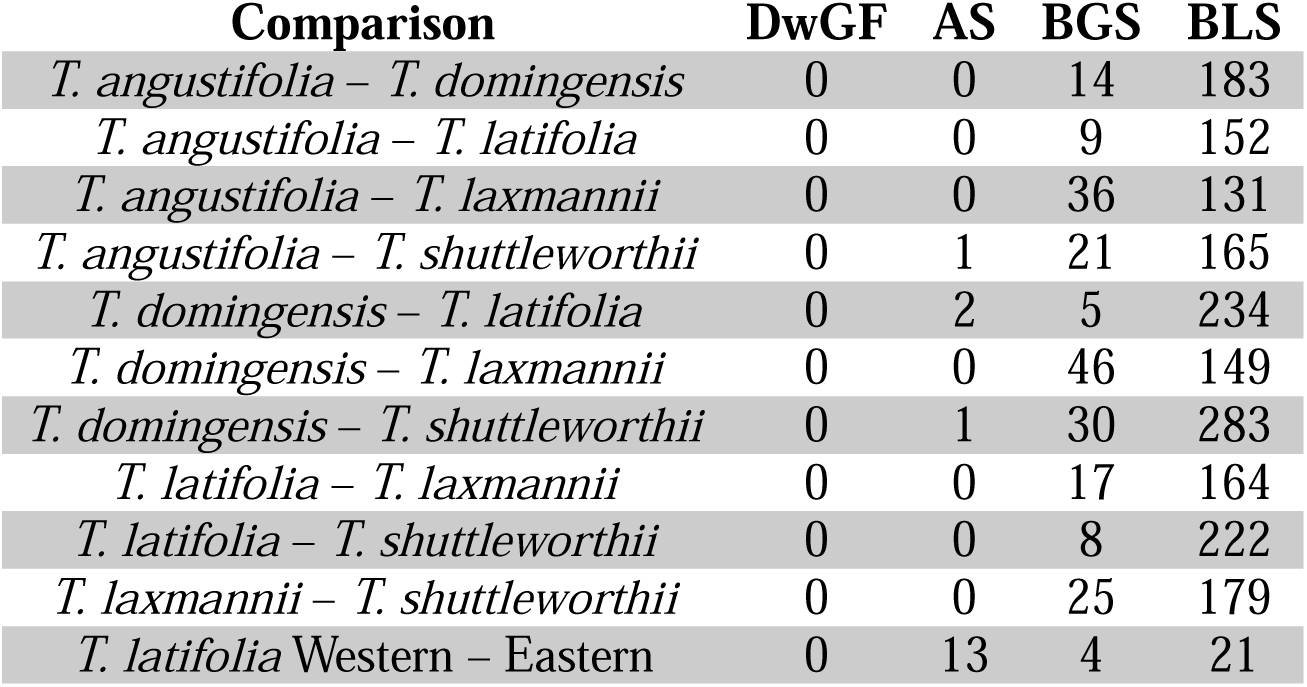
Pairwise genomic scan results. Total number of outlier windows (10 Kb each) under one of the four models of ecological divergence proposed by Han *et al*. (2017) and Irwin *et al*. (2018); (DwGF) divergent selection under gene flow, (AS) divergent selection without gene flow, (BGS) background selection, (BLS) balancing selection

| Comparison | DwGF | AS | BGS | BLS |
| --- | --- | --- | --- | --- |
| *T. angustifolia* – *T. domingensis* | 0 | 0 | 14 | 183 |
| *T. angustifolia* – *T. latifolia* | 0 | 0 | 9 | 152 |
| *T. angustifolia* – *T. laxmannii* | 0 | 0 | 36 | 131 |
| *T. angustifolia* – *T. shuttleworthii* | 0 | 1 | 21 | 165 |
| *T. domingensis* – *T. latifolia* | 0 | 2 | 5 | 234 |
| *T. domingensis* – *T. laxmannii* | 0 | 0 | 46 | 149 |
| *T. domingensis* – *T. shuttleworthii* | 0 | 1 | 30 | 283 |
| *T. latifolia* – *T. laxmannii* | 0 | 0 | 17 | 164 |
| *T. latifolia* – *T. shuttleworthii* | 0 | 0 | 8 | 222 |
| *T. laxmannii* – *T. shuttleworthii* | 0 | 0 | 25 | 179 |
| *T. latifolia* Western – Eastern | 0 | 13 | 4 | 21 |

## Discussion

We reconstructed the speciation history of five cattail species; our results suggest that the focal species in this study diverged under geographic isolation during periods roughly contemporaneous with historical bottlenecks and expansions in *Typha*. Aridity was widespread during the Mid-Miocene (∼16–11.6 Ma) and Pliocene (∼5.3 to 2.6 Ma) (Scotese *et al*., 2021), particularly across Asia (Sun *et al*., 2015; Herbert *et al*., 2016; Shen *et al*., 2018; Butiseacă *et al*., 2021; Ao *et al*., 2021), the centre of origin of *Typha* (Zhou *et al*., 2018). This aridity could have fragmented the extent and connectivity of wetlands, altering the distribution and abundance of *Typha* and resulting in small effective population sizes and increased genetic drift, thereby triggering allopatric diversification. Our results add to the evidence that climatically driven speciation was ubiquitous worldwide (Hewitt, 2000, 2001; Shafer *et al*., 2010; Kumar and Kumar, 2018).

For the demographic history of *T. angustifolia* relative to *T. domingensis*, we estimated population sizes, divergence and secondary contact times, and migration rates using a model based on the AIC-selected model but incorporating N_e_ shifts over time. Unmodelled size changes can lead to extreme biases in estimates of divergence times (Momigliano *et al*., 2021). This decision was supported by the congruence between the divergence time estimated by the chloroplast phylogeny (∼9.9 Ma) and that of the model we chose (∼9.22 Ma). Overall, modelling of species’ demographic histories suggested that divergence events occurred in isolation and were followed by secondary contact.

The genomic landscapes indicated the absence of divergent selection driving species differentiation. This result suggests that ecological speciation played a minor role in the divergence of *Typha* in this study. Some signatures of divergent selection may have been eroded over time, consistent with late-stage divergence (Burri *et al*., 2015), since *Typha* is an old genus—the age we estimated is 41.9 Ma, and Zhou *et al*. (2018) dated it at ∼39.03 Ma. However, the net divergence (d_a_) among species was relatively low (<0.02, i.e., within the “grey zone of speciation”, Roux *et al*., 2016), indicating weak reproductive isolation and thus rejecting the idea of late-stage divergence. Low and slowly accumulated differentiation is expected under a drift-driven divergence (reviewed in Turelli *et al*., 2001), and could be one explanation for some cattails’ ability to hybridise.

Genetic diversity reflects the reservoir of traits and potential responses to environmental changes that species encounter, thereby influencing their resilience across different ecosystems (Gregorius, 1987; Frankham *et al*., 2002; Hartl and Clark, 2006). Excluding *T. latifolia*, species’ genetic diversity was high (π ≥ 0.01; Begun *et al*., 2007), consistent with current large population sizes and substantial gene flow. Reduced genetic diversity is considered detrimental because it can reflect low adaptive potential (Teixeira and Huber, 2021; Kardos *et al*., 2021). However, some taxa, including invasive species, thrive despite low genetic diversity (Tsutsui *et al*., 2000; Roman and Darling, 2007; Charlesworth and Jensen, 2022). *Typha* can tolerate a wide range of climates, nutrients, pollutants, and pH and water levels (Sojda and Solberg, 1993; Kadlec and Wallace, 2008; Sesin *et al*., 2021), and the most widespread and commonly recognised cattail species is *T. latifolia* (Smith, 1987), known to have large census sizes and to be invasive in Oceania (Xu et al., 2013). Low diversity in *T. latifolia* suggests that alternative mechanisms, such as epigenetic modifications and phenotypic plasticity, may underlie this species’ success (Mounger *et al*., 2021). In *Arabidopsis thaliana*, epigenetic diversity underlies morphological variation and phenotypic plasticity when genetic diversity is reduced (Kooke et al., 2015; Schmid et al., 2018; Zhang et al., 2013). Future research could test whether epigenetic changes or phenotypic plasticity are more extensive in *T. latifolia* than in other *Typha* species.

## Conclusions

Understanding the causes of speciation is a central focus of evolutionary biology. We tested the roles of allopatry and selection in driving speciation in *Typha*, a globally distributed plant genus foundational to freshwater ecosystems. Our results suggest that bottlenecks and geographic isolation were drivers of speciation in some *Typha* lineages. The frequency of secondary contact with gene flow across species after drift-driven divergence (e.g., Pettengill and Moeller, 2012; Schield *et al*., 2019; Dong *et al*., 2020; Arteaga *et al*., 2020; Yamasaki *et al*., 2020; Le Provost *et al*., 2022; Wang *et al*., 2022; Kessler *et al*., 2023) raises the question of whether genetic barriers to gene flow are stronger or arise faster between lineages diverging under ecological speciation (e.g., Papadopulos *et al*., 2019; Liu *et al*., 2020; Christmas *et al*., 2021) than between those diverging in allopatry. Biologists now have access to novel data and methods for investigating this question (e.g., Laetsch *et al*., 2023; Burban *et al*., 2024), offering a complementary lens on the evolution of reproductive isolation.

## Acknowledgements

The laboratory procedures and data analyses in this study were conducted at Trent University, located on the traditional territory of the Mississauga Anishinaabeg, to whom we pay our respects. We thank Polina Volkova and Tulsi Patel for their invaluable contributions in the field and the laboratory. This work was funded by the Natural Sciences and Engineering Research Council of Canada, and Alberto Aleman was supported by the Environmental and Life Sciences Graduate Program at Trent University. SHARCNET and The Digital Research Alliance of Canada provided computational resources for this study. We are grateful to Camille Kessler, Marie-Laurence Cossette, the Associate Editors, and the anonymous reviewers for their valuable feedback, which greatly improved the quality of this manuscript. Finally, we thank Enrique Ruiz for his work on Figure 1 and Collin Wilson for his suggestions to improve the efficiency of ∂a∂i.

## Author Contribution Statement

Conceptualisation, developing methods, conducting research, data interpretation, writing, data analysis, preparation of figures and tables: Aaron Shafer, Alberto Aleman, Joanna Freeland, and Marcel Dorken.

All authors contributed to the manuscript and approved its final version.

## Data archiving

Upon acceptance of this article, we will submit the sequencing data to the Dryad data repository. Scripts and VCF files for this study are available at https://github.com/al-aleman/totoras_hdy and https://gitlab.com/WiDGeT_TrentU/graduate_theses/-/tree/master/aleman/hdy.

## Research Ethics Statement

This research did not require any ethical permissions.

