## Supplementary figures and images for "Repeated cases of allopatric divergence and secondary contact in cattail (*Typha*) speciation"

### Supplementary Figures

#### Slide 1
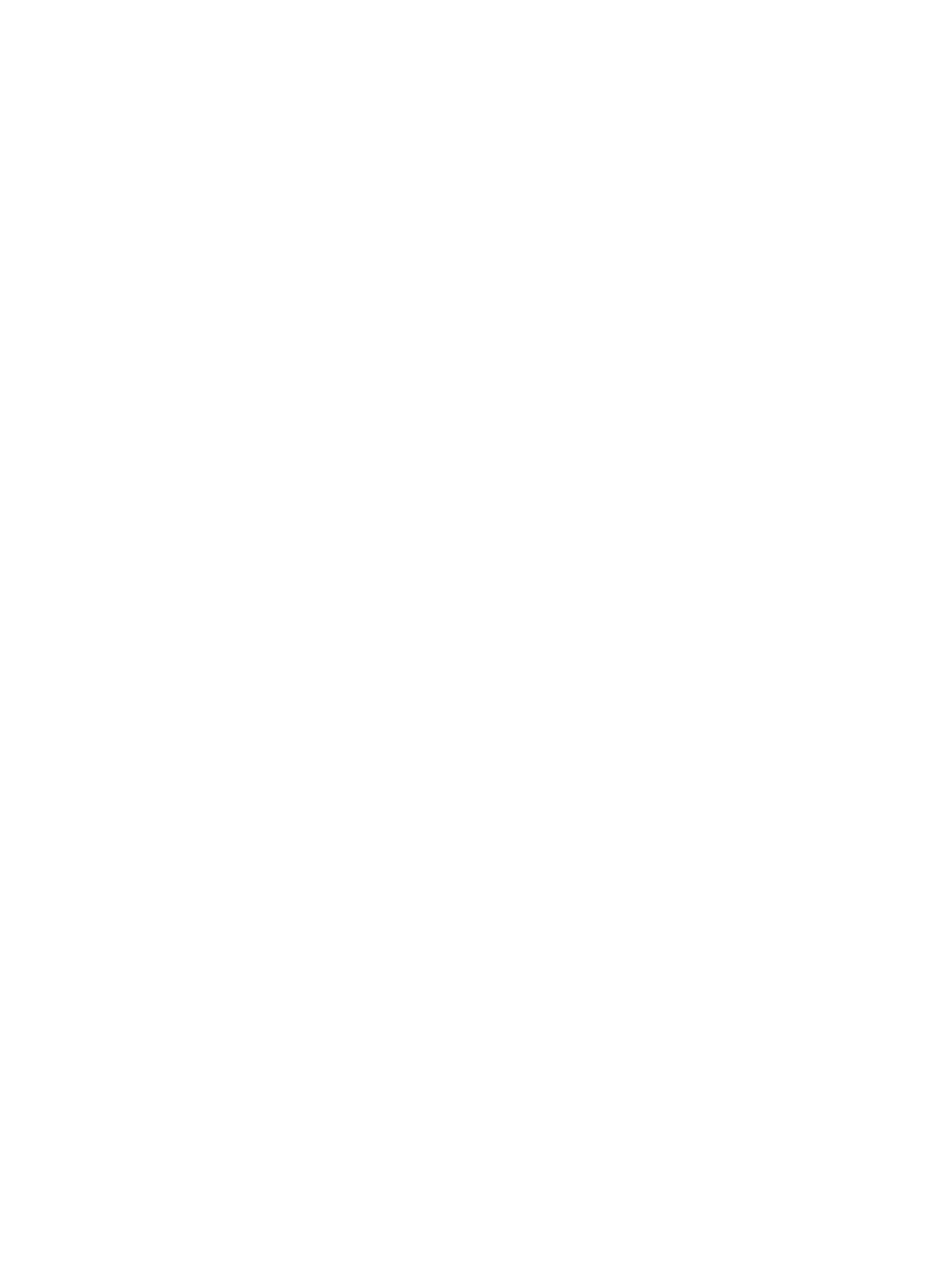

#### Slide 2
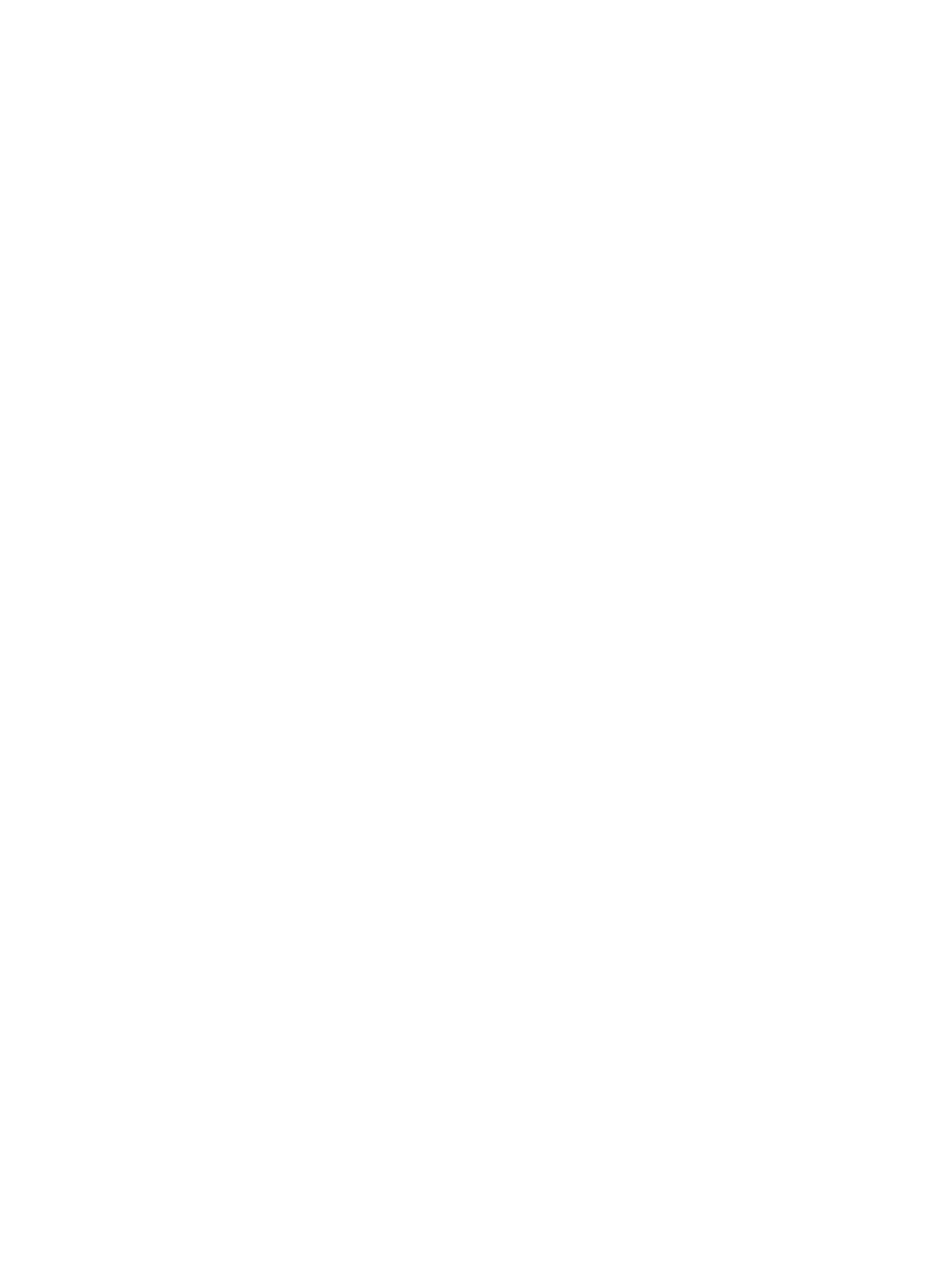

#### Slide 3
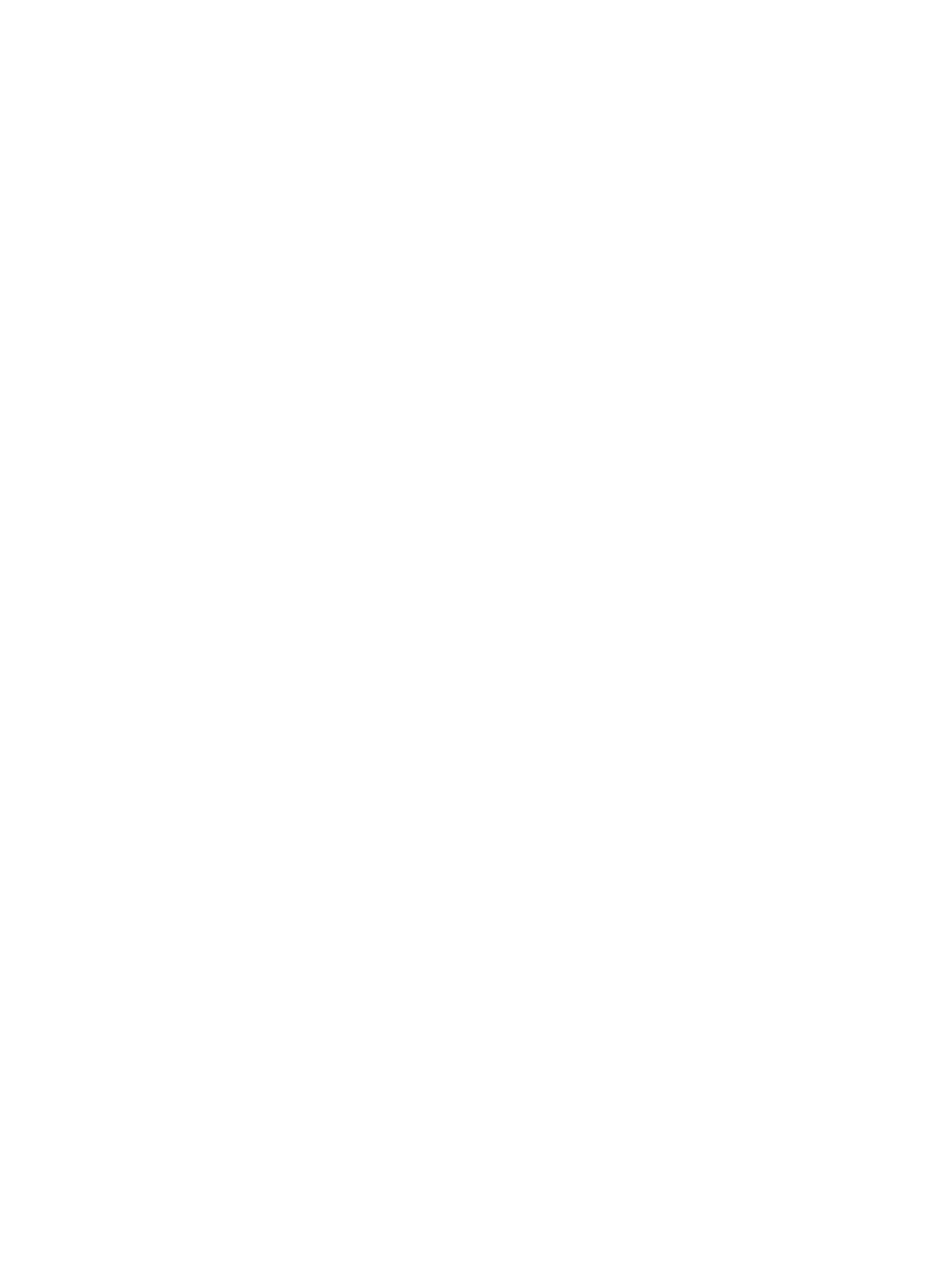
